# Asymmetrical template-DNA strand segregation can explain density-associated mutation-rate plasticity

**DOI:** 10.1101/307371

**Authors:** Duur K. Aanen, Alfons J.M. Debets

## Abstract

The mutation rate is a fundamental factor in evolutionary genetics. Recently, mutation rates were found to be strongly reduced at high density in a wide range of unicellular organisms, prokaryotic and eukaryotic. Independently, cell division was found to become more asymmetrical at increasing density in diverse organisms; in yeast, some ‘mother’ cells continue dividing, while their ‘offspring’ cells do not divide further. Here, we investigate how this increased asymmetry in cell division at high density can be reconciled with reduced mutation-rate estimates. We calculated the expected number of mutant cells due to replication errors under various modes of segregation of template-DNA strands and copy-DNA strands, both under exponential and under linear growth. We show that the observed reduction in the mutation rate at high density can be explained if mother cells preferentially retain the template-DNA strands, since new mutations are then confined to non-dividing daughter cells thus reducing the spread of mutant cells. Any other inheritance mode results in an *increase* in the number of mutant cells at higher density. The proposed hypothesis that patterns of DNA-strand segregation are density dependent fundamentally challenges our current understanding of mutation-rate estimates and extends the distinction between germline and soma to unicellular organisms.

## 1. Introduction

Mutation rates are typically minimized, as far as population genetic constraints allow [1]. However, mutation rates can vary, not only between organisms but also with environmental conditions. A recent study identifies a completely unexpected kind of mutation-rate plasticity in response to population density [2], which is dependent on quorum sensing [3]. Across a wide range of unicellular organisms, both eukaryotic and prokaryotic, the mutation rate consistently was found to decrease with increasing population density, with up to 23-fold lower mutation rates at high density than at low density. We propose a model that attributes reduced mutation rate at high density to increased asymmetry in mutation acquisition between ‘mother’ cells and ‘offspring’ cells, and discuss recent experimental studies that support this model.

It was long believed that unicellular organisms potentially do not age, thus exhibiting functional immortality. However, the last two decades have seen increasing evidence for asymmetrical cell division leading to differential cell fates, even in organisms with morphologically symmetrical division, such as *Escherichia coli* and fission yeast [4, 5]. An asymmetrical cell division results in a senescing ‘mother’ cell and a rejuvenated ‘daughter’ cell, and fecundity of the mother cell decreases with each division as damaged proteins and cell components accumulate. There is increasing evidence that such asymmetries during cell division are not limited to physiological and morphological cell characteristics, but extend to patterns of DNA strand inheritance, as shown in yeast [6, 7] and *E. coli* [8] and various types of stem cells [9].

The ‘Immortal Strand Hypothesis’ proposes that asymmetries in DNA-strand inheritance reduce the number of mutations in somatic cells [10]. According to this hypothesis, adult stem cells have ‘Template Strand Co-segregation’ (TSC [9, 10]), where the daughter cell maintaining the stem-cell function retains specific ‘master’ templates of the DNA strands of each chromosome (the parental strands [11]) at each division, while the differentiating daughter cell receives the new, ‘copy’ strands. Since most mutations during replication occur in the newly synthesized DNA strands and fewer in the template strands, this asymmetrical distribution reduces the mutation rate in the stem cells [10]. In support of the immortal strand hypothesis, TSC during cell division has been demonstrated in a broad range of organisms [9, 12-14], although it is not universal for stem cells and alternative hypotheses have been proposed to explain it [9, 15].

Recently, the degree of asymmetry during cell division was found to be higher at high density, in independent studies, for budding yeast [16] and for *E. coli* [17]. Furthermore, for muscle stem cells asymmetry of strand segregation was found to be increased when stem cells were seeded at higher cell densities [18]. Here we investigate how those findings of increased asymmetries at high density can be reconciled with reduced estimates of the mutation rate under that condition [2, 3]. We show that the observed reduction in the mutation rate at high density can be explained if mother cells preferentially retain the template-DNA strands, since new mutations are then confined to non-dividing daughter cells thus reducing the spread of mutant cells.

## 2. Methods

### Calculating the number of mutant cells resulting from a single mutational event

To establish the effect of the mode of DNA-strand segregation under different modes of cell division, we calculate the average number of mutant cells when a single copy error occurs during the formation of a certain number of cells from a single ancestor. For both symmetrical and asymmetrical cell division, we considered the effect of asymmetries in the distribution of template-DNA strands and copy-DNA strands between cells. Even though it seems unlikely that those asymmetries will be absolute, for the sake of argument, we consider deterministic models for both symmetrical and asymmetrical growth, under three different inheritance patterns of template DNA and copy-DNA strands by mother and daughter cells (Figure 1): (i) the copied strands are inherited by the mother cell (copy-strand co-segregation; CSC); (ii) the template strands are inherited by the mother cell (template-strand co-segregation; TSC); and (iii) the template and copied strands are inherited randomly between mother and daughter cells (random strand segregation; RSS). In all cases, we consider mutations due to copy error, the most common class of mutations [19], so mutations occur in the copied strand, and not in the template strand. We first show the calculation for the expected number of mutant cells for the formation of four cells from a single ancestor, and then for the general case.

**Figure 1.**
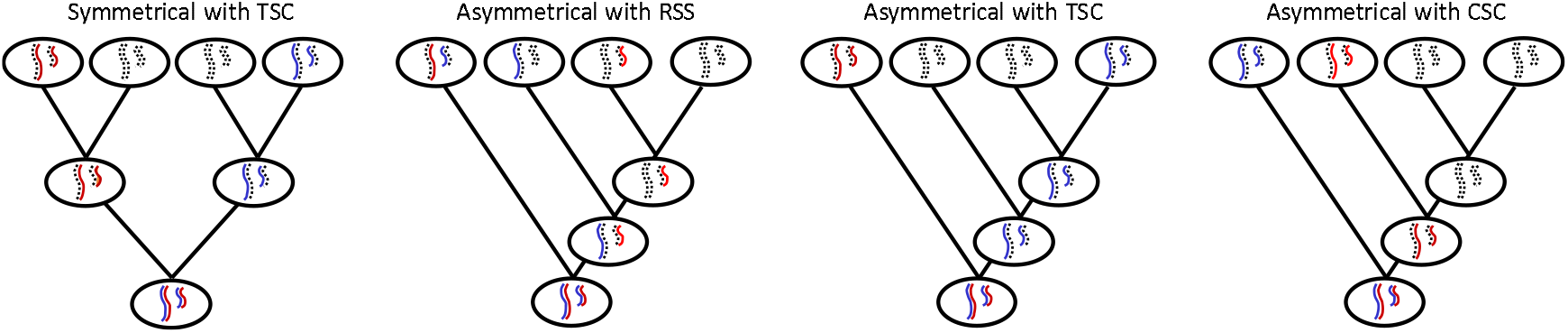
A comparison between symmetrical cell division with template strand co-segregation (TSC; left) and asymmetrical cell division with three forms of DNA-strand inheritance: RSS (center left), TSC (center right) and CSC (right). Following DNA replication, an asymmetrical cell division results in two daughters, one of which becomes a new mother, and the other of which is a rejuvenated cell that stops dividing. According to the ‘immortal strand hypothesis’ the sister chromatids containing the older strands (blue non-dashed) are retained in the continually dividing mother cell. Since the segregation pattern does not influence the number of mutant cells when cell division is symmetrical, symmetrical cell division is only drawn for one type of strand inheritance.

For symmetrical growth with RSS, the mutation can occur during the first division for which there are two routes or the second division, for which there are four routes. In the first case, two mutant cells will result, in the second case a single mutant cell. So we get 2+2+1+1+1+1=8 possibilities to get mutant cells and we have six routes for those to occur, so on average this yields 8/6=1.33 mutant cells. For symmetrical cell division with TSC or CSC, the expected number of mutant cells is the same. For *k* rounds of symmetrical division (with RSS, TSC or CSC), the number of mutant cells produced by a single mutational event can be calculated as: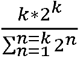.

For asymmetrical growth with RSS, a single mutation either ends up in a daughter cell (left branch), or in the mother cell (right branch). In the first case it will yield a single mutant cell only. If the mutation ends in the mother cell, it can either yield three mutant cells (when the mutation occurs in the first division), two mutant cells (when the mutation occurs in the second division) or a single mutant cell (when the mutation occurs in the third division). The average number of mutant cells if the mutation occurs in the mother cell is thus two. So the average number of mutant cells resulting from a single mutational event if cells division is asymmetrical and strands segregate randomly is (1+2)/2=1.5. More generally, for *n* cells formed with asymmetrical cell division, the expected number of mutant cells resulting from a single mutational event is 0.5+0.25*n*.

For asymmetrical growth with CSC, a single mutation always is inherited by the right branch. Depending on the timing, it can either yield three mutant cells (when the mutation occurs in the first division), two mutant cells (when the mutation occurs in the second division) or a single mutant cell (when the mutation occurs in the third division). The average number of mutant cells if the mutation occurs in the mother cell is thus two. More generally, for *n* cells formed with asymmetrical cell division, the expected number of mutant cells resulting from a single mutational event is 0.5*n*.

For asymmetrical cell division with TSC, the mutation will always end in a non-dividing daughter cell, and thus only yield a single mutant cell.

## 3. Results

Consider a culture of unicellular organisms grown at high nutrition (Figure 2). Initially, when nutrition is not limiting yet, growth will be maximal and exponential [20]. When nutrition becomes limiting, growth will increasingly become non-exponential (Figure 2a). As has been shown for yeast at high density, mother cells start to act like stem-cell lineages that continue budding off rejuvenated offspring cells for their entire replicative lifespan or for the remainder of it, while the rejuvenated offspring cells are quiescent and do not divide further [21] (Figure 2b). At low nutrition, the transition to non-exponential growth is less strictly associated with differentiation between mother cells and rejuvenated offspring cells [16-18].

**Figure 2.**
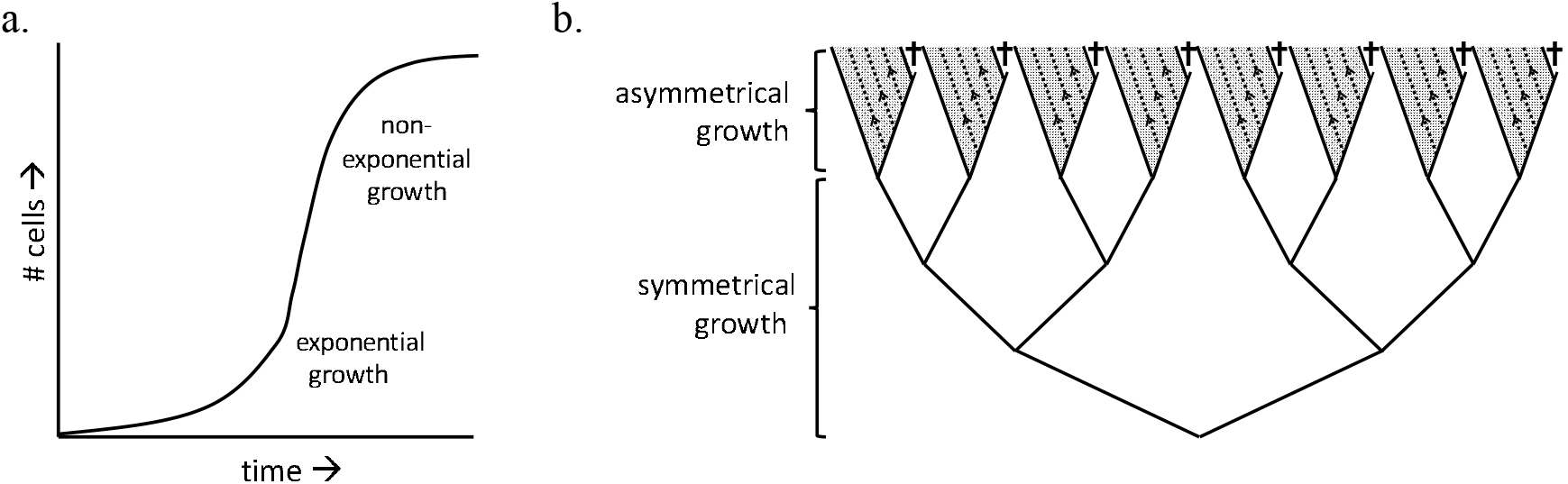
How growth changes from symmetrical exponential to asymmetrical and linear at high density. **a.** Initially, when nutrition is not limiting yet, exponential growth occurs. At higher density the culture is still growing, be it increasingly non-exponentially. **b.** Schematic representation of the shift from exponential to linear growth. At high density, cells increasingly start to divide asymmetrically. Senescing ‘mother’ cells act as stem-cell lineages, continuing to bud off ‘offspring’ cells, which stop dividing.

To explore the effect of growth mode on the expected number of mutant cells, we considered the number of mutant cells due to copy errors, which occur in the newly synthesised DNA strand. For symmetrical growth, this number does not depend on inheritance patterns of DNA template and copy strands, since all cells continue dividing. However, for asymmetrical growth, the number of mutant cells is influenced by the inheritance pattern of DNA strands. To establish this effect, we determined the expected number of mutant cells when a single copy error occurs during the formation of a certain number of cells, for both symmetrical and asymmetrical growth, under three different inheritance patterns of template and copy-strands by mother and daughter cells (Figure 1; Methods).

For asymmetrical growth, the inheritance pattern has a strong effect on the expected number of mutant cells. Consider asymmetrical division of a mother cell that can still bud off 31 daughter cells. CSC would then be expected to yield an average of no less than 16 mutant cells, RSS 8.5, and TSC only a single mutant cell (see Methods and Figure 3 for the calculation). The latter number is also lower than the number expected with symmetrical growth, where a single mutation in one of five subsequent rounds of divisions will yield an average of 2.6 mutant cells among the 32 resulting cells (irrespectively of the pattern of template and copy-strand segregation; Table I).

**Figure 3.**
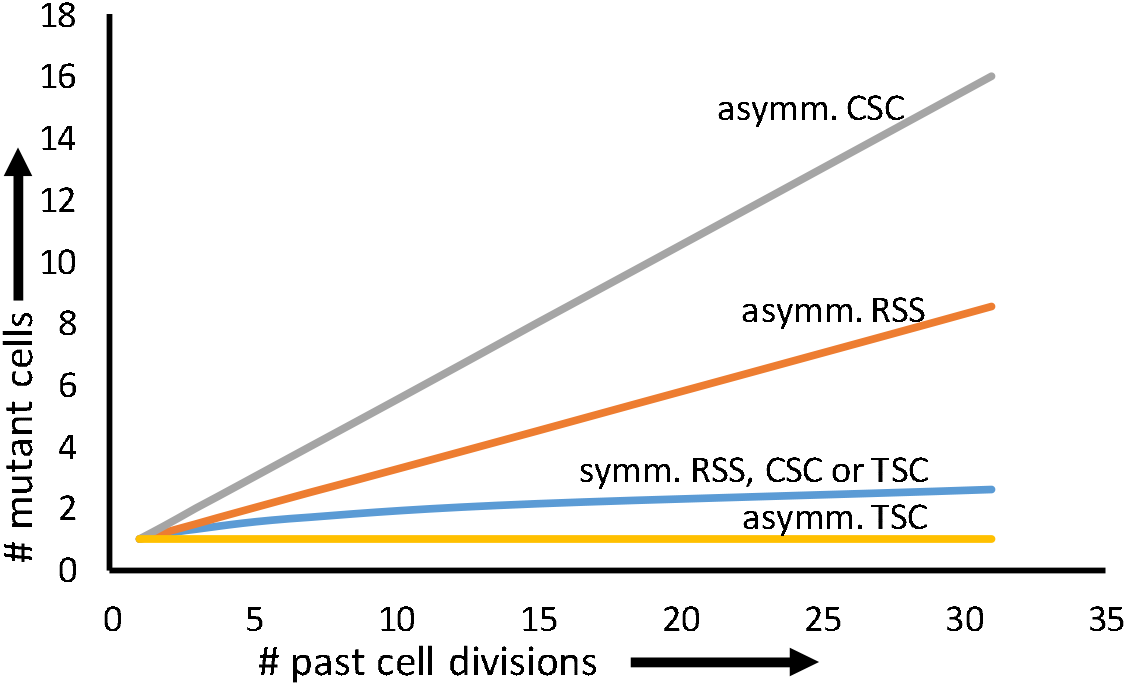
The expected number of mutant cells resulting from a single mutational event as a function of the number of past cell divisions under symmetrical cell division (with CSC, TSC or RSS; blue line; this number can be calculated as 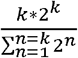 with *k* number of rounds of cell division with symmetrical growth), asymmetrical cell division with CSC (grey line; this number can be calculated as 0.5*n*, with *n* number of cells formed), asymmetrical cell division with RSS (orange line; this number can be calculated as 0.5+0.25*n*, with *n* number of cells formed) and asymmetrical division with TSC (yellow line; always 1, irrespectively of the number of cell divisions). The number of mutant cells under symmetrical division and asymmetrical division with RSS and CSC are all increasing functions of the number of cell divisions and all yield a higher number of mutant cells than asymmetrical division with TSC, which always yields only a single mutant cell, irrespective of the number of cell divisions.

**Table 1.**
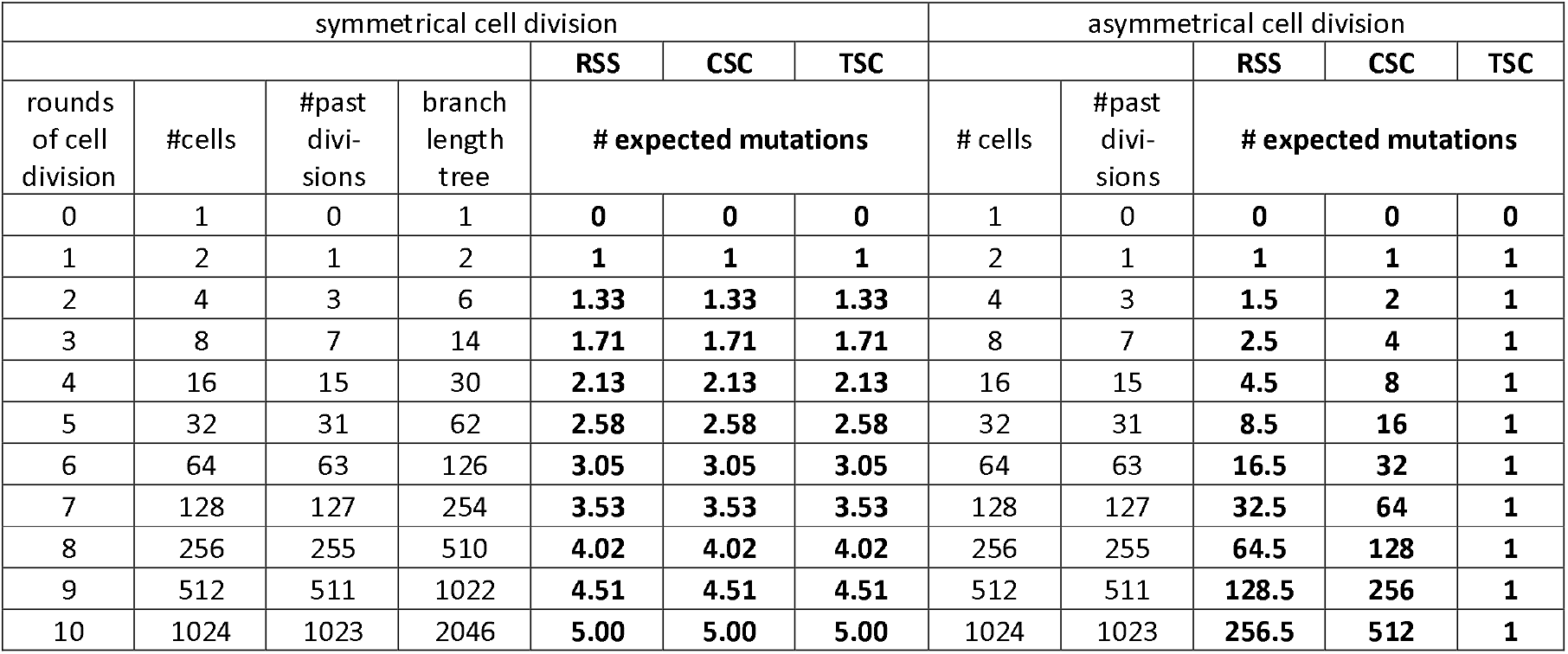
The expected numbers of mutant cells if one mutational event occurs during the formation of *n* cells under symmetrical and asymmetrical cell division, and with either RSS, CSC or TSC.

| symmetrical cell division |  |  |  |  |  |  | asymmetrical cell division |  |  |  |  |
| --- | --- | --- | --- | --- | --- | --- | --- | --- | --- | --- | --- |
|  |  |  |  | RSS | CSC | TSC |  |  | RSS | CSC | TSC |
| rounds of cell division | #cells | #past divisions | branch length tree | # expected mutations |  |  | # cells | #past divisions | # expected mutations |  |  |
| 0 | 1 | 0 | 1 | 0 | 0 | 0 | 1 | 0 | 0 | 0 | 0 |
| 1 | 2 | 1 | 2 | 1 | 1 | 1 | 2 | 1 | 1 | 1 | 1 |
| 2 | 4 | 3 | 6 | 1.33 | 1.33 | 1.33 | 4 | 3 | 1.5 | 2 | 1 |
| 3 | 8 | 7 | 14 | 1.71 | 1.71 | 1.71 | 8 | 7 | 2.5 | 4 | 1 |
| 4 | 16 | 15 | 30 | 2.13 | 2.13 | 2.13 | 16 | 15 | 4.5 | 8 | 1 |
| 5 | 32 | 31 | 62 | 2.58 | 2.58 | 2.58 | 32 | 31 | 8.5 | 16 | 1 |
| 6 | 64 | 63 | 126 | 3.05 | 3.05 | 3.05 | 64 | 63 | 16.5 | 32 | 1 |
| 7 | 128 | 127 | 254 | 3.53 | 3.53 | 3.53 | 128 | 127 | 32.5 | 64 | 1 |
| 8 | 256 | 255 | 510 | 4.02 | 4.02 | 4.02 | 256 | 255 | 64.5 | 128 | 1 |
| 9 | 512 | 511 | 1022 | 4.51 | 4.51 | 4.51 | 512 | 511 | 128.5 | 256 | 1 |
| 10 | 1024 | 1023 | 2046 | 5.00 | 5.00 | 5.00 | 1024 | 1023 | 256.5 | 512 | 1 |

In Table I and Figure 3, the expected number of mutant cells for a single mutational event as a function of the number of past cell division is given for the four possible combinations of division and strand segregation. As can be seen, for asymmetrical growth with TSC the expected number of mutant cells is one, irrespectively of the number of cell divisions. In contrast, for symmetrical growth, the expected number of mutant cells increases with the number of cell divisions and this is independent of the segregation mode of template and daughter strands. For asymmetrical growth with RSS and especially with CSC, the number of mutant cells increases even stronger with the number of cell divisions. For microorganisms where the maximal replicative lifespan has been determined at some 30 cell divisions [22], a 2.6-fold reduction in the expected number of mutant cells compared to symmetrical division leading to the same number of cells seems the maximum.

## 4. Discussion

Our results show that the empirical finding of a reduced mutation rate at high density can be reconciled with increased asymmetry in cell division under that condition if TSC occurs in the mother cells, which continue to divide. Asymmetrical cell division with TSC can account for a significant reduction in mutation-rate estimates, although not sufficient to fully explain the density-dependent mutation-rate plasticity recently reported [2, 3]. However, there is another catch when growth shifts from exponential to linear. Estimates of the mutation rate assume exponential growth [23]. The fluctuation test takes into account the probability that a mutation occurs at an early growth stage, in which case a large proportion of the population will have the mutation (a so-called ‘jackpot’). The model proposed here, with linear growth by division from a stem-cell mother that retains the template strands will never yield a ‘jackpot’, since mutations in the non-exponential phase always occur in terminal branches. This implies that the mutation rate will be systematically underestimated, which may account for the remaining difference. Furthermore, if asymmetrical growth occurs in the later stages of both low-density and high-density conditions, but TSC only at high density, the difference in mutation rate between low and high density will further increase (Figure 3).

The apparent universality of density-associated mutation-rate plasticity begs for a general mechanism. Given the independent evidence for a link between the degree of asymmetrical cell division and density in widely divergent organisms as bacteria [17], single-celled eukaryotes [16] and stem cells of multicellular eukaryotes [18], it seems plausible that this mechanism is based on asymmetrical cell division. The model proposed here is best supported for yeast. In yeast, at high density, a larger fraction of the cells becomes quiescent, being arrested in the G_0_ phase of the cell cycle [16], and consisting almost exclusively of rejuvenated quiescent daughter cells with a high capacity to grow when conditions improve [21]. The remaining cells are heterogeneous and show senescence. In support of a role for TSC, in yeast asymmetries in kinetochore inheritance have been shown [6], and one study found support for asymmetrical strand segregation [7], although another study did not [24]. However, the latter study used a low population density, which may account for this difference.

It seems paradoxical that the senescing cell retains the template DNA strands, and thus acquires the fewest mutations, while the rejuvenated offspring cells receive the copied strands, and thus any mutant cells. However, as explained above this inheritance pattern reduces the number of mutants among the rejuvenated cells. Perhaps the strongest argument in favour of the model proposed in this article is that the mutation rate will be strongly *increased* and not decreased if DNA strands were inherited randomly when cell division becomes asymmetrical. Even for the production of 16 rejuvenated cells by a mother cell, asymmetrical cell division with RSS would yield 4.5 times more mutant cells than asymmetrical cell division with TSC, and still over two times more than symmetrical division (Figure 3). The finding of on one hand, a reduction in the mutation rate at high density [2, 3] and on the other hand, an increase in asymmetrical division at high density [16-18], therefore makes it plausible that template-strand co-segregation occurs. However, direct evidence for our model remains to be provided. Recent improvements in the detection of mutations in single cells may make it feasible to test our hypothesis directly [25, 26]. An intriguing question is whether our model also applies to density-associated mutation rate plasticity found in viruses [2]. Since viruses are dependent on their host for genome replication, in the experiments used to measure the mutation rates at various density, virus density may correspond to host density, in which case our model may also apply to viral replication. It has been proposed that the mutation rate of RNA viruses may also depend on their replication mode, either by exponential replication where copy strands are copied or linear replication where template strands are used for replication only [27].

The plasticity in mutation rate in response to population density implies that numbers of mutational events per space and time vary much less with final population size than expected from a fixed mutation rate per cell division. In other words, the total number of cells with mutations occurring in a high-density and a low-density culture of unicellular organisms are more similar than expected based on the number of cell divisions that have occurred. This buffered number of mutant cells per space and time fits remarkably well in an emerging picture that the mutation rate of organisms is reduced by specific aspects of their growth mode, not only for vertebrate animals, which set aside germ cells early in development, but also for organisms that do not. For example, taller, long-lived plants have been found to have lower rates of molecular evolution per unit time than small plants, implying that the mutation rates per generation are more similar [28]. In plant meristems, the stem cells from which reproductive organs will develop undergo a minimal number of divisions during plant growth [29]. Also, the number of cell divisions separating axilliary meristems from the central meristem is minimized [30]. Similarly, in a fungus with an estimated age of more than 1,500 years the number of mutations was much lower than expected, presumably due to an unknown mechanism to reduce the number of mitotic divisions of cells at the growth front [31, 32]. In ciliates, a transcriptionally silent germline nucleus is present, whose mutation rate per cell division is more than an order of magnitude lower than that of other eukaryotes, but, converted to a per-sexual generation mutation rate, is remarkably similar to that of multicellular eukaryotes with a similar genome size [33].

The realisation that unicellular organisms also have mechanisms to reduce the mutation rate makes the germline-soma distinction more general than once believed. August Weismann was the first to distinguish an immortal germline from a disposable soma and argued that variations within individuals cannot be transmitted to the germline [34]. Leo Buss challenged Weismann’s doctrine, noticing that an early germline sequestration as seen in vertebrates is rare among multicellular organisms [35]. The recent findings discussed in this paper, however, revive part of Weismann’s doctrine. A picture emerges that germline sequestration is not limited to some animals, but also occurs in plants, fungi and even unicellular organisms, although the timing of sequestration may vary between organism groups and with ecological conditions such as population density.

## Acknowledgements

We thank Ben Auxier, Gerdien de Jong, Marc Maas and Vidyanand Nanjundiah for useful comments on this manuscript, and Piter Bijma, Hanna Kokko, and Arjan de Visser for useful comments on an earlier version of it. DKA was supported by the Netherlands Organisation for Scientific Research (VICI; NWO 86514007).

## Author contributions

DKA designed the study and wrote the manuscript with input from AJMD.

## References

1. Drake J.W., Charlesworth B., Charlesworth D., Crow J.F. 1998 Rates of spontaneous mutation. Genetics 148(4), 1667–1686.

2. Krasovec R., Richards H., Gifford D.R., Hatcher C., Faulkner K.J., Belavkin R.V., Channon A., Aston E., McBain A.J., Knight C.G. 2017 Spontaneous mutation rate is a plastic trait associated with population density across domains of life. Plos Biology 15(8). (doi:10.1371/journal.pbio.2002731).

3. Krasovec R., Belavkin R.V., Aston J.A.D., Channon A., Aston E., Rash B.M., Kadirvel M., Forbes S., Knight C.G. 2014 Mutation rate plasticity in rifampicin resistance depends on Escherichia coli cell-cell interactions. Nature Communications 5. (doi:10.1038/ncomms4742).

4. Barker M.G., Walmsley R.M. 1999 Replicative ageing in the fission yeast Schizosaccharomyces pombe. Yeast 15(14), 1511–1518. (doi:10.1002/(sici)1097-0061(199910)15:14<1511::aid-yea482>3.0.co;2-y).

5. Stewart E.J., Madden R., Paul G., Taddei F. 2005 Aging and death in an organism that reproduces by morphologically symmetric division. Plos Biology 3(2), 295–300. (doi:10.1371/journal.pbio.0030045).

6. Thorpe P.H., Bruno J., Rothstein R. 2009 Kinetochore asymmetry defines a single yeast lineage. Proceedings of the National Academy of Sciences of the United States of America 106(16), 6673–6678. (doi:10.1073/pnas.0811248106).

7. Williamson D.H., Fennell D.J. 1981 Non-random assortment of sister chromatids in yeast mitosis. In Molecular Genetics in Yeast, Alfred Benson Symposium (ed. von Wettstein D F.J., Kielland-Brandt M, Stenderup M), pp. 89 –107. Copenhagen, Munksgaard.

8. White M.A., Eykelenboom J.K., Lopez-Vernaza M.A., Wilson E., Leach D.R.F. 2008 Non-random segregation of sister chromosomes in Escherichia coli. Nature 455(7217), 1248–1250. (doi:10.1038/nature07282).

9. Charville G.W., Rando T.A. 2011 Stem cell ageing and non-random chromosome segregation. Philos Trans R Soc B-Biol Sci 366(1561), 85–93. (doi:10.1098/rstb.2010.0279).

10. Cairns J. 1975 Mutation selection and natural history of cancer. Nature 255(5505), 197–200. (doi:10.1038/255197a0).

11. Meselson M., Stahl F.W. 1958 The replication of DNA in Escherichia coli. Proceedings of the National Academy of Sciences 44(7), 671–682. (doi:10.1073/pnas.44.7.671).

12. Gurevich D.B., Nguyen P.D., Siegel A.L., Ehrlich O.V., Sonntag C., Phan J.M.N., Berger S., Ratnayake D., Hersey L., Berger J., et al. 2016 Asymmetric division of clonal muscle stem cells coordinates muscle regeneration in vivo. Science 353(6295), 136-+. (doi:10.1126/science.aad9969).

13. Rosenber.Rf, Kessel M. 1968 Nonrandom sister chromatid segregation and nuclear migration in hyphae of Aspergillus nidulans. Journal of Bacteriology 96(4), 1208-&.

14. Snedeker J., Wooten M., Chen X. 2017 The Inherent Asymmetry of DNA Replication. In Annual Review of Cell and Developmental Biology, Vol 33 (ed. Schekman R.), pp. 291–318.

15. Lansdorp P.M. 2007 Immortal strands? Give me a break. Cell 129(7), 1244–1247. (doi:10.1016/j.cell.2007.06.017).

16. Leonov A., Feldman R., Piano A., Arlia-Ciommo A., Lutchman V., Ahmadi M., Elsaser S., Fakim H., Heshmati-Moghaddam M., Hussain A., et al. 2017 Caloric restriction extends yeast chronological lifespan via a mechanism linking cellular aging to cell cycle regulation, maintenance of a quiescent state, entry into a non-quiescent state and survival in the non-quiescent state. Oncotarget 8(41), 69328–69350. (doi:10.18632/oncotarget.20614).

17. Lele U.N., Baig U.I., Watve M.G. 2011 Phenotypic Plasticity and Effects of Selection on Cell Division Symmetry in Escherichia coli. Plos One 6(1). (doi:10.1371/journal.pone.0014516).

18. Shinin V., Gayraud-Morel B., Gomes D., Tajbakhsh S. 2006 Asymmetric division and cosegregation of template DNA strands in adult muscle satellite cells. Nature Cell Biology 8(7), 677–U669. (doi:10.1038/ncb1425).

19. Crow J.F. 2000 The origins patterns and implications of human spontaneous mutation. Nature Reviews Genetics 1(1), 40–47. (doi:10.1038/35049558).

20. Monod J. 1949 The growth of bacterial cultures. Annual Review of Microbiology 3, 371–394. (doi:10.1146/annurev.mi.03.100149.002103).

21. Allen C., Buttner S., Aragon A.D., Thomas J.A., Meirelles O., Jaetao J.E., Benn D., Ruby S.W., Veenhuis M., Madeo F., et al. 2006 Isolation of quiescent and nonquiescent cells from yeast stationary-phase cultures. Journal of Cell Biology 174(1), 89–100. (doi:10.1083/jcb.200604072).

22. Nystrom T., Liu B.D. 2014 The mystery of aging and rejuvenation - a budding topic. Current Opinion in Microbiology 18, 61–67. (doi:10.1016/j.mib.2014.02.003).

23. Luria S.E., Delbrück M. 1943 Mutations of bacteria from virus sensitivity to virus resistance. Genetics 28(6), 491–511.

24. Neff M.W., Burke D.J. 1991 Random segregation of chromatids at mitosis in Saccharomyces cerevisiae. Genetics 127(3), 463–473.

25. Elez M., Murray A.W., Bi L.J., Zhang X.E., Matic I., Radman M. 2010 Seeing Mutations in Living Cells. Current Biology 20(16), 1432–1437. (doi:10.1016/j.cub.2010.06.071).

26. Robert L., Ollion J., Robert J., Song X.H., Matic I., Elez M. 2018 Mutation dynamics and fitness effects followed in single cells. Science 359(6381), 1283–1286. (doi:10.1126/science.aan0797).

27. Sardanyes J., Martinez F., Daros J.A., Elena S.F. 2012 Dynamics of alternative modes of RNA replication for positive-sense RNA viruses. Journal of the Royal Society Interface 9(69), 768–776. (doi:10.1098/rsif.2011.0471).

28. Lanfear R., Ho S.Y.W., Davies T.J., Moles A.T., Aarssen L., Swenson N.G., Warman L., Zanne A.E., Allen A.P. 2013 Taller plants have lower rates of molecular evolution. Nature Communications 4. (doi:10.1038/ncomms2836).

29. Heidstra R., Sabatini S. 2014 Plant and animal stem cells: similar yet different. Nature Reviews Molecular Cell Biology 15(5), 301–312. (doi:10.1038/nrm3790).

30. Burian A., de Reuille P.B., Kuhlemeier C. 2016 Patterns of Stem Cell Divisions Contribute to Plant Longevity. Current Biology 26(11), 1385–1394. (doi:10.1016/j.cub.2016.03.067).

31. Aanen D.K. 2014 DEVELOPMENTAL BIOLOGY How a long-lived fungus keeps mutations in check. Science 346(6212), 922–923. (doi:10.1126/science.1261401).

32. Anderson J.B., Catona S. 2014 Genomewide mutation dynamic within a long-lived individual of Armillaria gallica. Mycologia 106(4), 642–648.

33. Sung W., Tucker A.E., Doak T.G., Choi E., Thomas W.K., Lynch M. 2012 Extraordinary genome stability in the ciliate Paramecium tetraurelia. Proceedings of the National Academy of Sciences of the United States of America 109(47), 19339–19344. (doi:10.1073/pnas.1210663109).

34. Weismann A. 1893 The germ-plasm: a theory of heredity. New York, Charles Scribner’s sons.

35. Buss L.W. 1983 Evolution, development and the units of selection. Proceedings of the National Academy of Sciences of the United States of America-Biological Sciences 80(5), 1387–1391. (doi:10.1073/pnas.80.5.1387).

